# Modelling personality, plasticity and predictability in shelter dogs

**DOI:** 10.1101/145367

**Authors:** Conor Goold, Ruth C. Newberry

**Affiliations:** Department of Animal and Aquacultural Sciences, Faculty of Biosciences, Norwegian University of Life Sciences

**Keywords:** inter- and intra-individual differences, behavioural reaction norms, behavioural repeatability, longitudinal behavioural assessment, human-animal interactions

## Abstract

Behavioural assessments of shelter dogs (*Canis lupus familiaris*) typically comprise standardised test batteries conducted at one time point but test batteries have shown inconsistent predictive validity. Longitudinal behavioural assessments offer an alternative. We modelled longitudinal observational data on shelter dog behaviour using the framework of behavioural reaction norms, partitioning variance into personality (i.e. inter-individual differences in behaviour), plasticity (i.e. individual differences in behavioural change) and predictability (i.e. individual differences in residual intra-individual variation). We analysed data on 3,263 dogs’ interactions (N = 19,281) with unfamiliar people during their first month after arrival at the shelter. Accounting for personality, plasticity (linear and quadratic trends) and predictability improved the predictive accuracy of the analyses compared to models quantifying personality and/or plasticity only. While dogs were, on average, highly sociable with unfamiliar people and sociability increased over days since arrival, group averages were unrepresentative of all dogs and predictions made at the individual level entailed considerable uncertainty. Effects of demographic variables (e.g. age) on personality, plasticity and predictability were observed. Behavioural repeatability was higher one week after arrival compared to arrival day. Our results highlight the value of longitudinal assessments on shelter dogs and identify measures that could improve the predictive validity of behavioural assessments in shelters.

## Introduction

*Personality*, defined by inter-individual differences in average behaviour, represents just one component of behavioural variation of interest in animal behaviour research. Personality frequently describes less than 50% of behavioural variation in animal personality studies [1, 2], leading to the combined analysis of personality with *plasticity*, individual differences in behavioural change [3], and *predictability*, individual differences in residual intra-individual variability [4–8]. Understanding these different sources of behavioural variation simultaneously can be achieved using the general framework of behavioural reaction norms [3, 5], which provides insight into how animals react to fluctuating environments through time and across contexts. The concept of behavioural reactions norms is built upon the use of hierarchical statistical models to quantify between- and within-individual variation in behaviour, following methods in quantitative genetics [3]. More generally, these developments reflect increasing interest across biology in expanding the ‘trait space’ of phenotypic evolution [9] beyond mean trait differences and systematic plasticity across environmental gradients to include residual trait variation (e.g. developmental instability: [10, 11]; stochastic variation in gene expression: [12]).

Modest repeatability of behaviour has been documented in domestic dogs (*Canis lupus familiaris*), providing evidence for personality variation. For instance, using meta-analysis, Fratkin *et al.* [13] found an average Pearson's correlation of behaviour through time of 0.43, explaining 19% of the behavioural variance between successive time points (where the average time interval between measurements was 21 weeks). However, the goal of personality assessments in dogs is often to predict an individual dog's future behaviour (e.g. working dogs: [14, 15]; pet dogs: [16]) and, thus, it is important not to confuse the stability of an individual's behaviour relative to the behaviour of others with stability of intra-individual behaviour. That is, individuals could vary their behaviour in meaningful ways in response to internal (e.g. ontogeny) and external (e.g. environmental) factors while maintaining differences from other individuals. When time-related change in dog behaviour has been taken into account, behavioural change at the group-level has been of primary focus (e.g. [16–18]) and no studies have explored the heterogeneity of residual variance within each dog. The predominant focus on inter-individual differences and group-level patterns of behavioural change risks obscuring important individual-level heterogeneity and may partly explain why a number of dog personality assessment tools have been unreliable in predicting future behaviour [14–16,19].

Of particular concern is the low predictive value of shelter dog assessments for predicting behaviour post-adoption [20–24], resulting in calls for longitudinal, observational models of assessment [20, 24]. Animal shelters are dynamic environments and, for most dogs, instigate an immediate threat to homeostasis as evidenced by heightened hypothalamic-pituitary-adrenal axis activity and an increase in stress-related behaviours (e.g. [25–28]). Over time, physiological and behavioural responses are amenable to change [17,27,29]. Therefore, dogs in shelters may exhibit substantial heterogeneity in intra-individual behaviour captured neither by standardised behavioural assessments conducted at one time point [24] nor by group-level patterns of behavioural change. An additional complication is that the behaviour in shelters may not be representative of behaviour outside of shelters. For example, Patronek and Bradley [29] suggested that up to 50% of instances of aggression expressed while at a shelter are likely to be false positives. Such false positives may be captured in estimates of predictability, with individuals departing more from their representative behaviour having higher residual intra-individual variability (lower predictability) than others. Overall, absolute values of behaviour, such as mean trait values across time (i.e. personality), may account for just part of the important behavioural variation needed to understand and predict shelter dog behaviour. While observational models of assessment have been encouraged, methods to systematically analyse longitudinal data collected at shelters into meaningful formats are lacking.

In this paper, we demonstrate how the framework of behavioural reaction norms can be used to quantify inter- and intra-individual differences in shelter dog behaviour. To do so, we employ data on dogs' interactions with unfamiliar people from a longitudinal and observational shelter assessment. As a core feature of personality assessments, how shelter dogs interact with unknown people is of great importance. At one extreme, if dogs bite or attempt to bite unfamiliar people, they are at risk of euthanasia [29]. At the other extreme, even subtle differences in how dogs interact with potential adopters can influence adoption success [30]. Importantly, neither may all dogs react to unfamiliar people in the same way through time at the shelter nor may all dogs show the same day-to-day fluctuation of behaviour around their average behavioural trajectories. These considerations can be explored by examining behavioural reaction norms.

The analysis of behavioural reaction norms is dependent on the use of hierarchical statistical models for partitioning variance among individuals [3,5,6]. Given that ordinal data are common in behavioural research, here, we illustrate how similar hierarchical models can be applied to ordinal data using a Bayesian framework (see also [31]). Apart from distinguishing inter-from intra-individual variation, we place particular emphasis on two desirable properties of the hierarchical modelling approach taken here. First, the property of *hierarchical shrinkage* [32] offers an efficacious way of making inferences about individual-level behaviour when data are highly unbalanced and potentially unrepresentative of a dog's typical behaviour. When data are sparse for certain individuals, hierarchical shrinkage means that an individual's parameter estimates (e.g. intercepts) are more similar to, or shrunken, towards the group-level estimates. Secondly, since any prediction of future (dog) behaviour will entail uncertainty, a Bayesian approach is attractive because we can directly obtain a probability distribution of parameter values consistent with the data (i.e. the posterior distribution) for all parameters [32, 33]. By contrast, frequentist confidence intervals are not posterior probability distributions and, thus, their interpretation is more challenging when a goal is to understand uncertainty in parameter estimates [32].

## Material & Methods

### Subjects

Behavioural data on *N* = 3,263 dogs from Battersea Dogs and Cats Home's longitudinal, observational assessment model were used for analysis. The data concerned all behavioural records of dogs at the shelter during 2014 (including those arriving in 2013 or departing in 2015), filtered to include all dogs: 1) at least 4 months of age (to ensure all dogs were treated similarly under shelter protocols, e.g. vaccinated so eligible for walks outside and kennelled in similar areas), 2) with at least one observation during the first 31 days since arrival at the shelter, and 3) with complete data for demographic variables to be included in the formal analysis (Table 1). Because dogs spent approximately one month at the shelter on average (Table 1), we focused on this period in our analyses (arrival day 0 to day 30). We did not include breed characterisation due to the unreliability of using appearance to attribute breed type to shelter dogs of uncertain heritage [34].

**Table 1.** Demographic variables of dogs in the sample analysed. Mean and standard deviation (SD) or the number of dogs by category (N) are displayed.

| **Demographic variable** | **Mean (SD) / N** |
| --- | --- |
| Number of observations per dog | 5.9 (3.7) |
| Days spent at the shelter | 25.8 (35.0) |
| Age (years; all at least 4 months old) | 3.7 (3.0) |
| Weight (kg) | 18.9 (10.2) |
| Source: gift / stray / return | 1950 / 1122 / 191 |
| Rehoming centre: London / Old Windsor / Brands Hatch | 1873 / 951 / 439 |
| Females / males | 1396 / 1867 |
| Neutered: before arrival / at shelter / not / undetermined | 1043 / 1281 / 747 / 192 |

### Shelter environment

Details of the shelter environment have been presented elsewhere [35]. Briefly, the shelter was composed of three different rehoming centres (Table 1): one large inner-city centre based in London (approximate capacity: 150-200 dogs), a medium-sized suburban/rural centre based in Old Windsor (approximate capacity: 100-150 dogs), and a smaller rural centre in Brands Hatch (approximate capacity: 50 dogs). Dogs considered suitable for adoption were housed in indoor kennels (typically about 4m × 2m, with a shelf and bedding alcove; see also [36]). Most dogs were housed individually, and given daily access to an indoor run behind their kennel. Feeding, exercising and kennel cleaning were performed by a relatively stable group of staff members. Dogs received water ad libitum and two meals daily according to veterinary recommendations. Sensory variety was introduced daily (e.g. toys, essential oils, classical music, access to quiet ‘chill-out’ rooms). Regular work hours were from 0800 h to 1700 h each day, with public visitation from 1000 h to 1600 h. Dogs were socialised with staff and/or volunteers daily.

### Data collection

The observational assessment implemented at the shelter included observations of dogs by trained shelter employees in different, everyday contexts, each with its own qualitative ethogram of possible behaviours. Shortly after dogs were observed in relevant contexts, employees entered observations into a custom, online platform using computers located in different housing areas. Each behaviour within a context had its own code. Previously, we have reported on aggressive behaviour across contexts [35]. Here, we focus on variation in behaviour in one of the most important contexts, ‘Interactions with unfamiliar people’, which pertained to how dogs reacted when people with whom they had never interacted before approached, made eye contact, spoke to and/or attempted to make physical contact with them. For the most part, this context occurred outside of the kennel, but it could also occur if an unfamiliar person entered the kennel. Observations could be recorded by an employee meeting an unfamiliar dog, or by an employee observing a dog meeting an unfamiliar person. Different employees could input records for the same dog, and employees could discuss the best code to describe a certain observation if required.

Behavioural observations in the ‘Interactions with unfamiliar people’ context were recorded using a 13-code ethogram (Table 2). Each behavioural code was subjectively labelled and generally defined, providing a balance between behavioural rating and behavioural coding methodologies. The ethogram represented a scale of behavioural problem severity and assumed adoptability (higher codes indicating higher severity of problematic behaviour/lower sociability), reflected by grouping the 13 codes further into green, amber and red codes (Table 2). Green behaviours posed no problems for adoption, amber behaviours suggested dogs may require some training to facilitate successful adoption but did not pose a danger to people or other dogs, and red behaviours suggested dogs needed training or behavioural modification to facilitate successful adoption and could pose a risk to people or other dogs. A dog's suitability for adoption was, however, based on multiple behavioural observations over a number of days. When registering an observation, the employee selected the highest code in the ethogram that was observed on that occasion (i.e. the most severe level of problematic behaviour was given priority). There were periods when a dog could receive no entries for the context for several days but other times when multiple observations were recorded on the same day, usually when a previous observation was followed by a more serious behavioural event. In these instances, and in keeping with the shelter protocol, we retained the highest (i.e. most severe) behavioural code registered for the context that day. When the behaviours were the same, only one record was retained for that day. This resulted in an average of 5.9 (SD = 3.7; range = 1 to 22) records per dog on responses during interactions with unfamiliar people while at the shelter. For dogs with more than one record, the average number of days between records was 2.8 (SD = 2.2; range = 1 to 29).

**Table 2.** Ethogram of behavioural codes used to record observations of interactions with unfamiliar people, and their percent prevalence in the sample. Behaviour labels followed by + indicate a more intense form of the behaviour with the same name without a +.

| **Behaviour** | **Colour** | **%** | **Definition** |
| --- | --- | --- | --- |
| 1: Friendly | Green | 63.5 | Dog initiates interactions with people in an appropriate social manner. |
| 2: Excitable | Green | 14.2 | Animated interaction with an enthusiastic attitude, showing behaviours such as jumping up, mouthing, an inability to stand still, and/or playful behaviour towards people. |
| 3: Independent | Green | 4.1 | Does not actively seek interaction, although relaxed in the presence of people |
| 4: Submissive | Green | 4.6 | Appeasing and/or nervous behaviours, including a low body posture, rolling over and other calming signals. |
| 5: Uncomfortable avoids | Amber | 5.4 | Tense and stiff posture, and/or shows anxious behaviours (e.g. displacement behaviours) while trying to move away from the person. |
| 6: Submissive + | Amber | 0.2 | High intensity of submissive behaviours such as submissive urination, a reluctance to move, or is frequently overwhelmed by the interaction. |
| 7: Uncomfortable static | Amber | 0.8 | Tense and stiff posture, and/or shows anxious behaviour (potentially showing displacement behaviours) but doesn't move away from the person. |
| 8: Stressed | Amber | 0.5 | High frequency/intensity of stress behaviours, which may include dribbling, stereotypic behaviours, stress vocalisations, constant shedding, trembling, and destructive behaviours. |
| 9: Reacts to people non-aggressive | Amber | 2.4 | Barks, whines, howls and/or play growls when seeing/meeting people, potentially pulling or lunging towards them. |
| 10: Uncomfortable approaches | Amber | 0.7 | Tense and stiff posture, and/or shows anxious behaviour (potentially showing displacement behaviours) and approaches the person. |
| 11: Overstimulated | Red | 0.8 | High intensity of excitable behaviour, including grabbing, body barging, and nipping. |
| 12: Uncomfortable static + | Red | 0.1 | Body freezes (the body goes suddenly and completely still) in response to an interaction with a person. |
| 13: Reacts to people aggressive | Red | 2.8 | Growls, snarls, shows teeth and/or snaps when seeing/meeting people, potentially pulling or lunging towards them. |

### Validity & inter-rater reliability

Inter-rater reliability and the validity of the assessment methodology were evaluated using data from a larger research project at the shelter. Videos depicting different behaviours in different contexts were filmed by canine behaviourists working at the shelter, who subsequently organised video coding sessions with 93 staff members (each session with about 5 - 10 participants) across rehoming centres [35]. The authors were blind to the videos and administration of video coding sessions. The staff members were shown 14 videos (each about 30 s long) depicting randomly-selected behaviours, two from each of seven different assessment contexts (presented in a pseudo-random order, the same for all participants). Directly after watching each video, they individually recorded (on a paper response form) which ethogram code best described the behaviour observed in each context. Two videos depicted behaviour during interactions with people (familiar versus unfamiliar not differentiated), one demonstrating *Reacts to people aggressive* and the other *Reacts to people non-aggressive* (Table 2). Below, we present the inter-rater reliabilities and the percentage of people who chose the correct behaviour and colour category for these two videos in particular, but also the averaged results across the 14 videos, since there was some redundancy between ethogram scales across contexts.

### Statistical analyses

All data analysis was conducted in R version 3.3.2 [37].

#### Validity & inter-rater reliability

Validity was assessed by calculating the percentage of people answering with the correct ethogram code/code colour for each video. Inter-rater reliability was calculated for each video using the consensus statistic [38] in the R package *agrmt* [39], which is based on Shannon entropy and assesses the amount of agreement in ordered categorical responses. A value of 0 implies complete disagreement (i.e. responses equally split between the lowest and highest ordinal categories, respectively) and a value of 1 indicates complete agreement (i.e. all responses in a single category). For the consensus statistic, 95% confidence intervals (CIs) were obtained using 10,000 non-parametric bootstrap samples. The confidence intervals were subsequently compared to 95% CIs of 10,000 bootstrap sample statistics from a null uniform distribution, which was created by: 1) selecting the range of unique answers given for a particular video and 2) taking 10,000 samples of the same size as the real data, where each answer had equal probability of being chosen. Thus, the null distribution represented a population with a realistic range of answers, but had no clear consensus about which category best described the behaviour. When the null and real consensus statistics' 95% CIs did not overlap, we inferred statistically significant consensus among participants.

#### Hierarchical Bayesian ordinal probit model

The distribution of ethogram categories was heavily skewed in favour of the green codes (Table 2), particularly the first *Friendly* category. Since some categories were chosen particularly infrequently, we aggregated the raw responses into a 6-category scale: 1) *Friendly*, 2) *Excitable*, 3) *Independent*, 4) *Submissive*, 5) *Amber codes*, 6) *Red codes*. This aggregated scale retained the main variation in the data and simplified the data interpretation. We analysed the data using a Bayesian ordinal probit model (described in [32, 40]), but extended to integrate the hierarchical structure of the data, including heteroscedastic residual standard deviations, to quantify predictability for each dog (for related models, see [31,41,42]). The ordinal probit model, also known as the cumulative or thresholded normal model, is motivated by a latent variable interpretation of the ordinal scale. That is, an ordinal dependent variable, *Y*, with categories *K_j_*, from *j* = 1 to *J*, is a realisation of an underlying continuous variable divided into thresholds, *θ*_*c*_,, for *C* = 1 to *J* - 1 Under the probit model, the probability of each ordinal category is equal to its area under the cumulative normal distribution, *ϕ*, with mean, *μ*, SD σ and thresholds *θ*_*c*_:

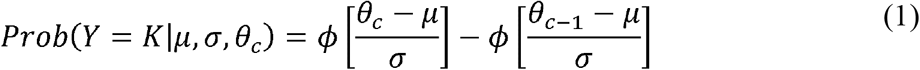

For the first and last categories, this simplifies to *ϕ* [(*θ*_c_ - *μ*) / σ] and 1 - *ϕ* [(*θ*_*c* - 1_ - *μ*) / σ], respectively. As such, the latent scale extends from.±∞ Here, the ordinal dependent variable was a realisation of the hypothesised continuum of ‘insociability when meeting unfamiliar people’, with 6 categories and 5 threshold parameters. While ordinal regression models usually fix the mean and SD of the latent scale to 0 and 1 and estimate the threshold parameters, we fixed the first and last thresholds to 1.5 and 5.5 respectively, allowing for the remaining thresholds, and the mean and SD, to be estimated from the data. As explained by Kruschke [32], this allows for the results to be interpretable with respect to the ordinal scale. We present the results using both the predicted probabilities of ordinal sociability codes and estimates on the latent, unobserved scale assumed to generate the ordinal responses.

#### Hierarchical structure

To model inter- and intra-individual variation, a hierarchical structure for both the mean and SD was specified. That is, parameters were included for both group-level and dog-level effects. The mean model, describing the predicted pattern of behaviour across days on the latent scale, *y*^*^, for observation *i* from dog *j*, was modelled as:

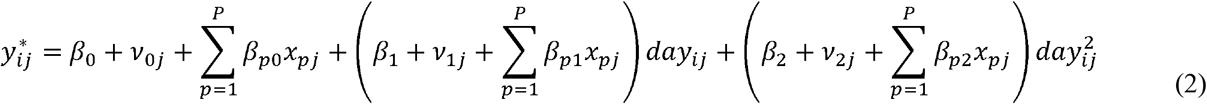

Equation 2 expresses the longitudinal pattern of behaviour as a function of i) a group-level intercept the same for all dogs, *β*_o_, and the deviation from the group-level intercept for each dog, *v*_0*j*_, ii) a linear effect of day since arrival, *β*_1_, and each dog's deviation, *v*_1*j*_, and iii) a quadratic effect of day since arrival, *β*_2_, and each dog's deviation, *v*_2*j*_. A quadratic effect was chosen based on preliminary plots of the data at group-level and at the individual-level, although we also compared the model's predictive accuracy with simpler models (described below). Day since arrival was standardised, meaning that the intercepts reflected the behaviour on the average day since arrival across dogs (approximately day 8). The three dog-level parameters, *v_j_*, correspond to personality and linear and quadratic plasticity parameters, respectively. The terms 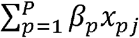 denote the effect of dog-level predictor variables (*x_p_*), included to explain variance between dog-level intercepts and slopes. These included: the number of observations for each dog, the number of days dogs spent at the shelter controlling for the number of observations (i.e. the residuals from a linear regression of total number of days spent at the shelter on the number of observations), average age while at the shelter, average weight at the shelter, sex, neuter status, source type, and rehoming centre (Table 1). For neuter status, we did not make comparisons between the ‘undetermined’ category and other categories. The primary goal of including these predictor variables was to obtain estimates of individual differences conditional on relevant inter-individual differences variables, since the data were observational.

The SD model was:

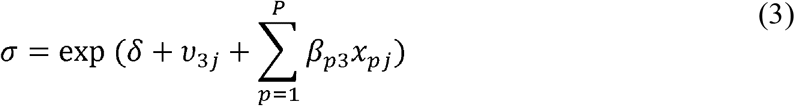

This equation models the SD of the latent scale by its own regression, with group-level SD intercept, *δ*, evaluated at the average day, the deviation for each dog from the group-level SD intercept, *v*_3*j*_, and predictor variables, 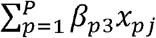, as in the mean model (equation 2). The SDs across dogs were assumed to approximately follow a log-normal distribution, with *l_n_*(σ)approximately normally distributed (hence the exponential inverse-link function). The parameter *v*_3*j*_ corresponds to each dog's residual SD or predictability.

All four dog-level parameters were assumed to be multivariate normally distributed with means 0 and variance-covariance matrix **Σ_v_** estimated from the data:

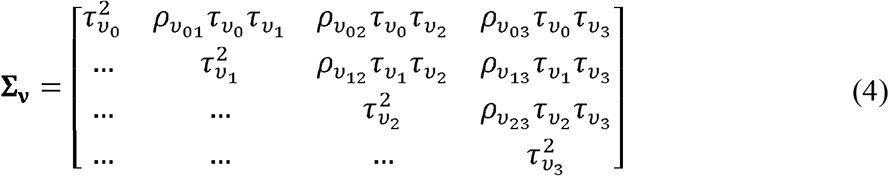

The diagonal elements are the variances of the dog-level intercepts, linear slopes, quadratic slopes and residual SDs, respectively, while the covariances fill the off-diagonal elements (only the upper triangle shown), where *ρ* is the correlation coefficient. In the results, we report *τ*_*v*3_ (the SD of dog-level residual SDs) on the original scale, rather than the log-transformed scale, using 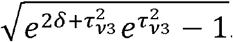 Likewise, *δ* was transformed to the median of the original scale by *e*^*δ*^.

To summarise the amount of behavioural variation explained by differences between individuals, referred to as repeatability in the personality literature [1], we calculated the intra-class correlation coefficient (ICC). Since the model includes both intercepts and slopes varying by dog, the ICC is a function of both linear and quadratic effects of day since arrival. The ICC for day i, assuming individuals with the same residual variance (i.e. using the median of the log-normal residual SD), was calculated as:

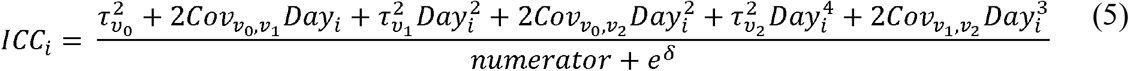

Equation 5 is an extension of the intra-class correlation calculated from mixed-effect models with a random intercept only [43] to include the variance parameters for, and covariances between, the linear and quadratic effects of day, which were evaluated at specific days of interest. We calculated the ICC for values of −1, 0 and 1 on the standardised day scale, corresponding to approximately the arrival day (day 0), day 8, and day 15. This provided a representative spread of days for most of the dogs in the sample, since there were fewer data available for later days which could lead to inflation of inter-individual differences.

To inspect the degree of rank-order change in sociability across dogs from arrival day compared to specific later days (i.e. whether dogs that were, on average, least sociable on arrival also tended to be least sociable later on), we calculated the ‘cross-environmental’ correlations [44] between the same days as the ICC. The cross-environmental covariance matrix, **Ω**, between the three focal days was calculated as:

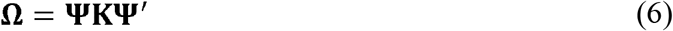

In equation 6, **Κ** represents the variance-covariance matrix of the dog-level intercepts and (linear and quadratic) slopes, and **Ψ** is a three-by-three matrix with a column vector of 1s, a column vector containing −1, 0, and 1 defining the day values for the cross-environmental correlations for the linear component, and a column vector containing 1, 0, and 1 defining the day values for the cross-environmental correlations for the quadratic component. Once defined, **Ω** was scaled to a correlation matrix. Finally, to summarise the degree of individual differences in predictability, we calculated the ‘coefficient of variation for predictability’ as 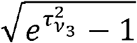 following Cleasby *et al.* [5].

#### Prior distributions

We chose prior distributions that were either weakly informative (i.e. specified a realistic range of parameter values) for computational efficiency, or weakly regularising to prioritise conservative inference. The prior for the overall intercept, *β*_0_, was *Normal*(y,5) where y is the arithmetic mean of the ordinal data. The linear and quadratic slope parameters, *β*_1_ and *β*_2_, were given *Normal* (0,1) priors. Coefficients for the dog-level predictor variables, *β_k_*, were given *Normal* (0,*σ_β_p__*) where *σ_β_p__* was a shared SD across predictor variables, which had in turn a half-Cauchy hyperprior with mode 0 and shape parameter 2, *half – Cauchy*(0,2). Using shared SD imposes shrinkage on the regression coefficients for conservative inference: when most regression coefficients are near zero, then estimates for other regression coefficients are also pulled towards zero (e.g. [32]). The prior for the overall log-transformed residual SD, *δ*, was *Normal* (0,1). The covariance matrix of the random effects was parameterised as a Cholesky decomposition of the correlation matrix (see [45] for more details), where the SDs had *half – Cauchy* (0,2).priors and the correlation matrix had a LKJ prior distribution [46] with shape parameter *η* set to 2.

### Model selection & computation

We compared the full model explained above to five simpler models. Starting with the full model, the alternative models included: i) parameters quantifying personality and quadratic and linear plasticity only; ii) parameters quantifying personality and linear plasticity only, with a fixed quadratic effect of day since arrival; iii) parameters quantifying personality only, with fixed linear and quadratic effects of day since arrival; iv) parameters quantifying personality only, with a fixed linear effect of day since arrival; and v) a generalised linear regression with no dog-varying parameters and a linear fixed effect for day since arrival (Figure 1). Models were compared by calculating the widely applicable information criterion (WAIC; [47]) following McElreath [33] (see the R script file). The WAIC is a fully Bayesian information criterion that indicates a model's *out-of-sample* predictive accuracy relative to other plausible models while accounting for model complexity, and is preferable to the deviance information criterion (DIC) because WAIC does not assume multivariate normality in the posterior distribution and returns a probability distribution rather than a point estimate [33]. Thus, WAIC guards against both under- and over-fitting to the data (unlike measures of purely in-sample fit, e.g. *R*^2^).

**Figure 1.**
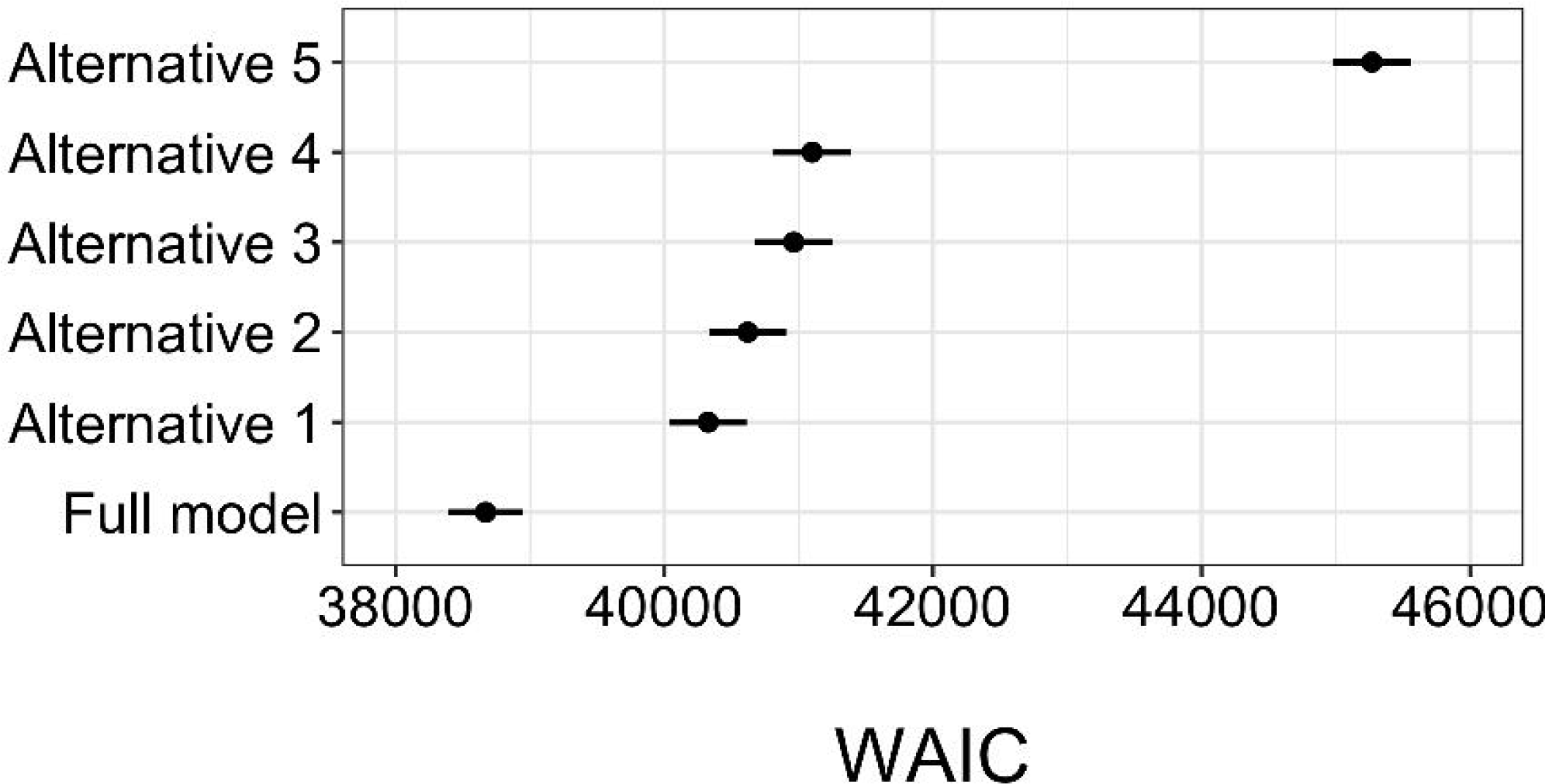
Out-of-sample predictive accuracy (lower is better) for each model (described in text section section 2.5.5) measured by the widely applicable information criterion (WAIC). Black points denote the WAIC estimate and horizontal lines show WAIC estimates ± standard error. Mean ± standard error: full model = 38669 ± 275; alternative 1 = 40326 ± 288; alternative 2 = 40621 ± 288; alternative 3 = 40963 ± 289; alternative 4 = 41100 ± 289; alternative 5 = 45268 ± 289.

Models were computed using the probabilistic programming language Stan [45] using the *RStan* package [48] version 2.15.1, which employs Markov chain Monte Carlo estimation using Hamiltonian Monte Carlo (see the R script file and Stan code for full details). We ran four chains of 5,000 iterations each, discarding the first 2,500 iterations of each chain as warm-up, and setting thinning to 1. Convergence was assessed visually using trace plots to ensure chains were well mixed, numerically using the Gelman-Rubin statistic (values close to 1 and < 1.05 indicating convergence) and by inspecting the effective sample size of each parameter. We also used graphical posterior predictive checks to assess model predictions against the raw data, including ‘counterfactual’ predictions [33] to inspect how dogs would be predicted to behave across the first month of being in the shelter regardless of their actual number of observations or length of stay at the shelter.

To summarise parameter values, we calculated mean (denoted *β*) and 95% highest density intervals (HDIs), the 95% most probable values for each parameter (using functions in the *rethinking* package; [33]). For comparing levels of categorical variables, the 95% HDI of their differences were calculated (i.e. the differences between the coefficients at each step in the MCMC chain, denoted *β_diff_*). When the 95% HDI of predictor variables surpassed zero, a credible effect was inferred.

## Results

### Inter-rater reliability & validity

For the two videos depicting interactions with people, consensus was 0.75 (95% CI: 0.66, 0.84) for the video showing an example of *Reacts to people non-aggressive* and 0.77 (95% CI: 0.74, 0.81) for the example of *Reacts to people aggressive*, respectively. Neither did these results overlap with the null distributions (see Supplementary Material Table S1), indicating significant inter-rater reliability. For the video showing *Reacts to people non-aggressive*, 77% chose the correct code and 83% a code of the correct colour category (amber), and, as previously reported by [35], 52% chose the correct code for the video showing *Reacts to people aggressive* and 55% chose a code of the correct colour category (red; 42% chose the amber code *Reacts to people non-aggressive* instead). Across all assessment context videos, the average consensus was 0.71 and participants chose the correct ethogram category 66% of the time while 78% of answers were a category of the correct ethogram colour.

### Hierarchical ordinal probit model

The full model had the best out-of-sample predictive accuracy, with the inclusion of heterogeneous residual SDs among dogs improving model fit by over 1,500 WAIC points compared to the second most plausible model (Alternative 1 in Figure 1). In general, models that included more parameters to describe personality, plasticity and predictability, and models with a quadratic effect of day, had better out-of-sample predictive accuracy, despite the added complexity brought by additional parameters.

At the group-level, the *Friendly* code (Table 2) was most probable overall and was estimated to increase in probability across days since arrival, while the remaining sociability codes either decreased or stayed at low probabilities (Figure 2a), reflecting the raw data. On the latent sociability scale (Figure 2b), the group-level intercept parameter on the average day was 0.68 (95% HDI: 0.51, 0.86). A one SD increase in the number of days since arrival was associated with a −0.63 unit (95% HDI: −0.77, −0.50) change on the latent scale on average (i.e. reflecting increasing sociability), and the group-level quadratic slope was positive (*β* = 0.20, 95% HDI: 0.10, 0.30), reflecting a quicker rate of change in sociability earlier after arrival to the shelter than later (i.e. a concave down parabola). There was a slight increase in the quadratic curve towards the end of the one-month period, although there were fewer behavioural observations at this point and so greater uncertainty about the exact shape of the curve, resulting in estimates being pulled closer to those of the intercepts. The group-level residual standard deviation had a median of 1.84 (95% HDI: 1.67, 2.02).

**Figure 2.**
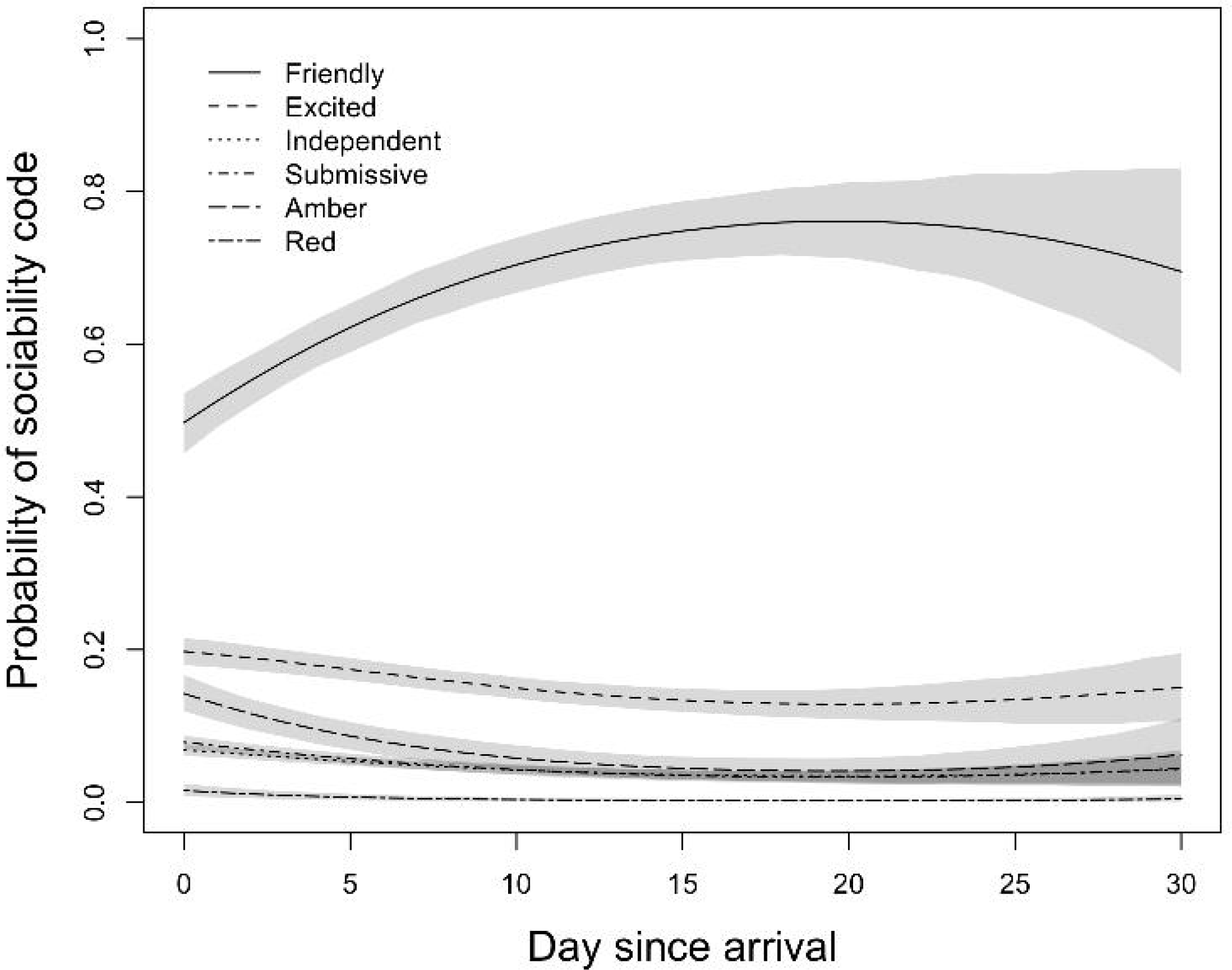
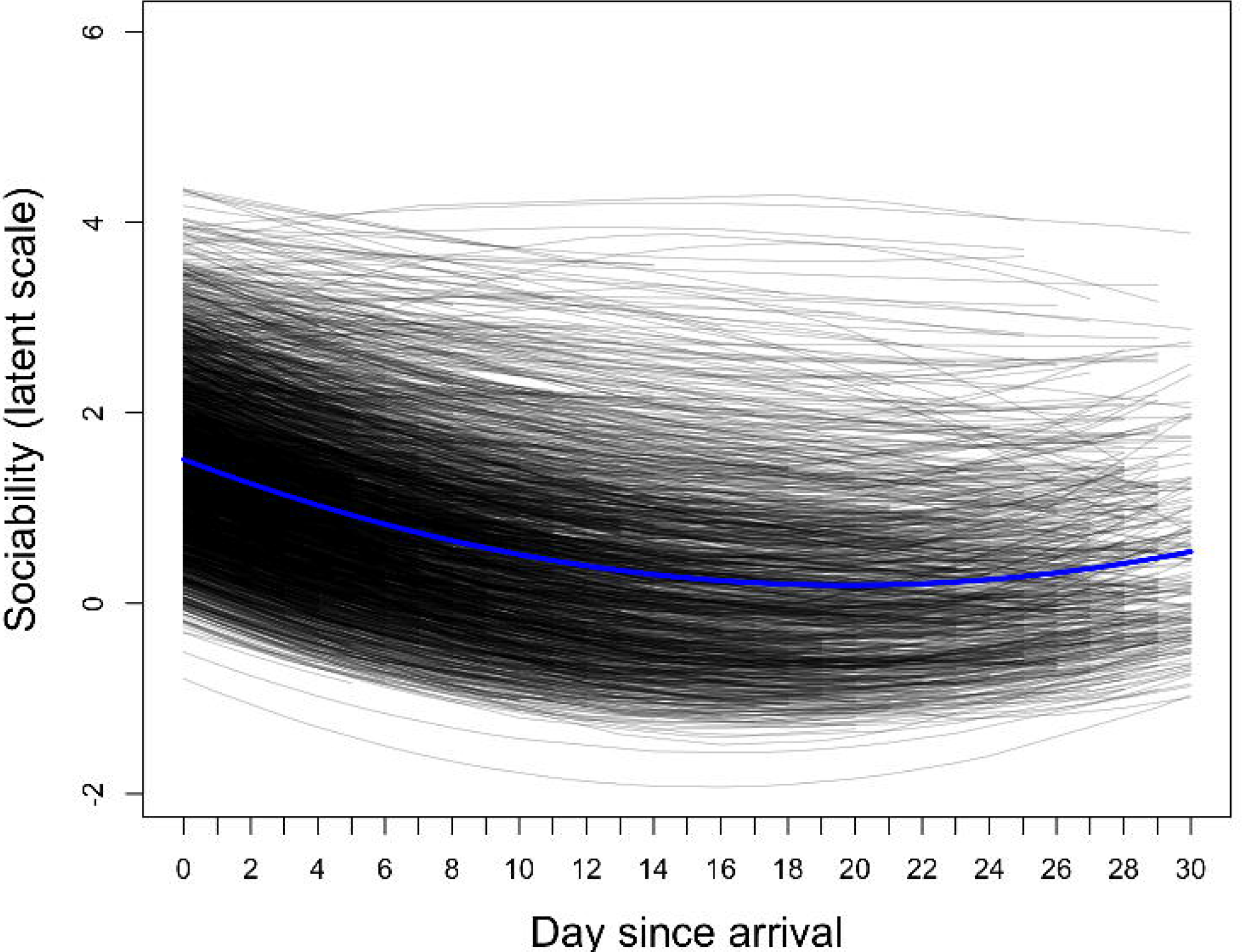
(a) Predicted probabilities (posterior means = black lines; 95% highest density intervals = shaded areas) of different sociability codes across days since arrival. (b) Posterior mean behavioural trajectories on the latent scale (ranging from± ∞) at the group-level (blue line) and for each individual (black lines), where higher values indicate lower sociability.

At the individual level, heterogeneity existed in behavioural trajectories across days since arrival (Figure 2b). The SDs of dog-varying parameters were: i) intercepts: 1.29 (95% HDI: 1.18, 1.41; Figure 3a), ii) linear slopes: 0.56 (95% HDI: 0.47, 0.65; Figure 3b), iii) quadratic slopes: 0.28 (95% HDI: 0.20, 0.35; Figure 3c), and iv) residual SDs: 1.39 (95% HDI: 1.22, 1.58; Figure 3d). There was also large uncertainty in individual-level estimates. Figure 4 displays counterfactual model predictions for twenty randomly-sampled dogs. Uncertainty in reaction norm estimates, illustrated by the width of the 95% HDIs (dashed black lines), was greatest when data were sparse (e.g. towards the end of the one-month study period). Hierarchical shrinkage meant that individuals with observations of less sociable responses, or individuals with few behavioural observations, tended to have model predictions pulled towards the overall mean. Note that regression lines depict values on the latent scale predicted to generate observations on the ordinal scale, and so may not clearly fit the ordinal data points. The coefficient of variation for predictability was 0.64 (95% HDI: 0.58, 0.70). Individuals with the five highest and lowest residual SD estimates are shown in Figure 5.

**Figure 3.**
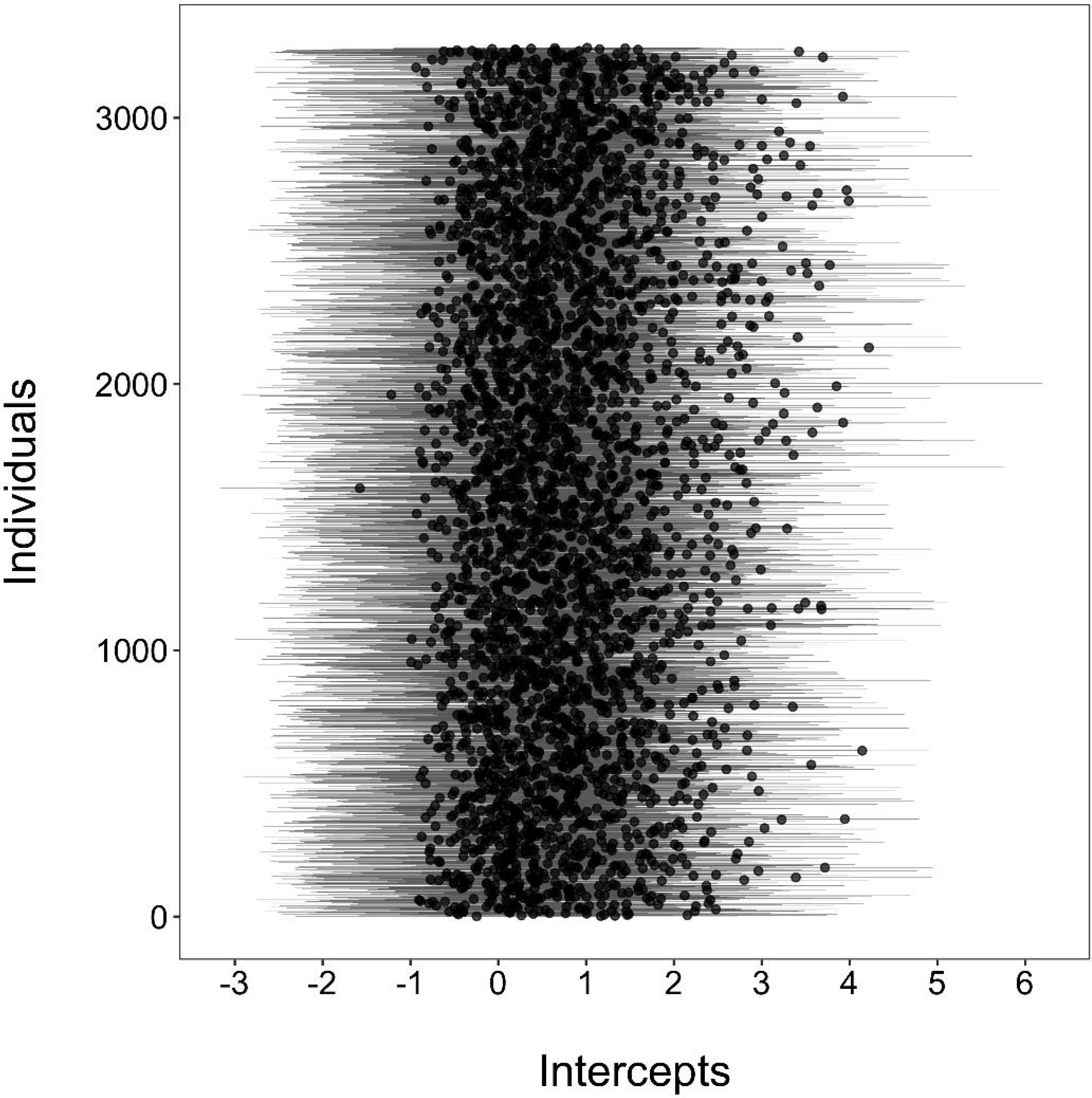
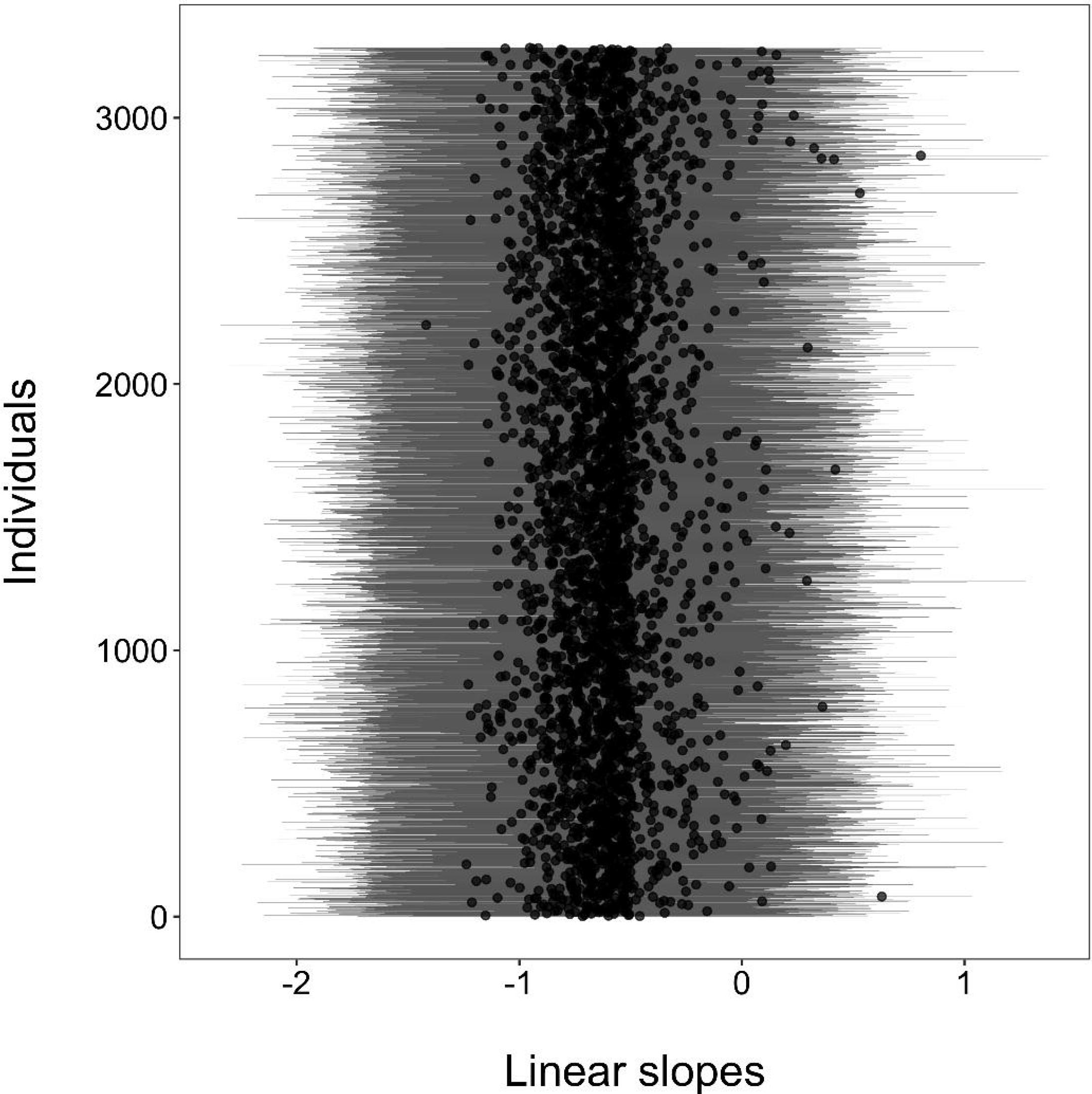
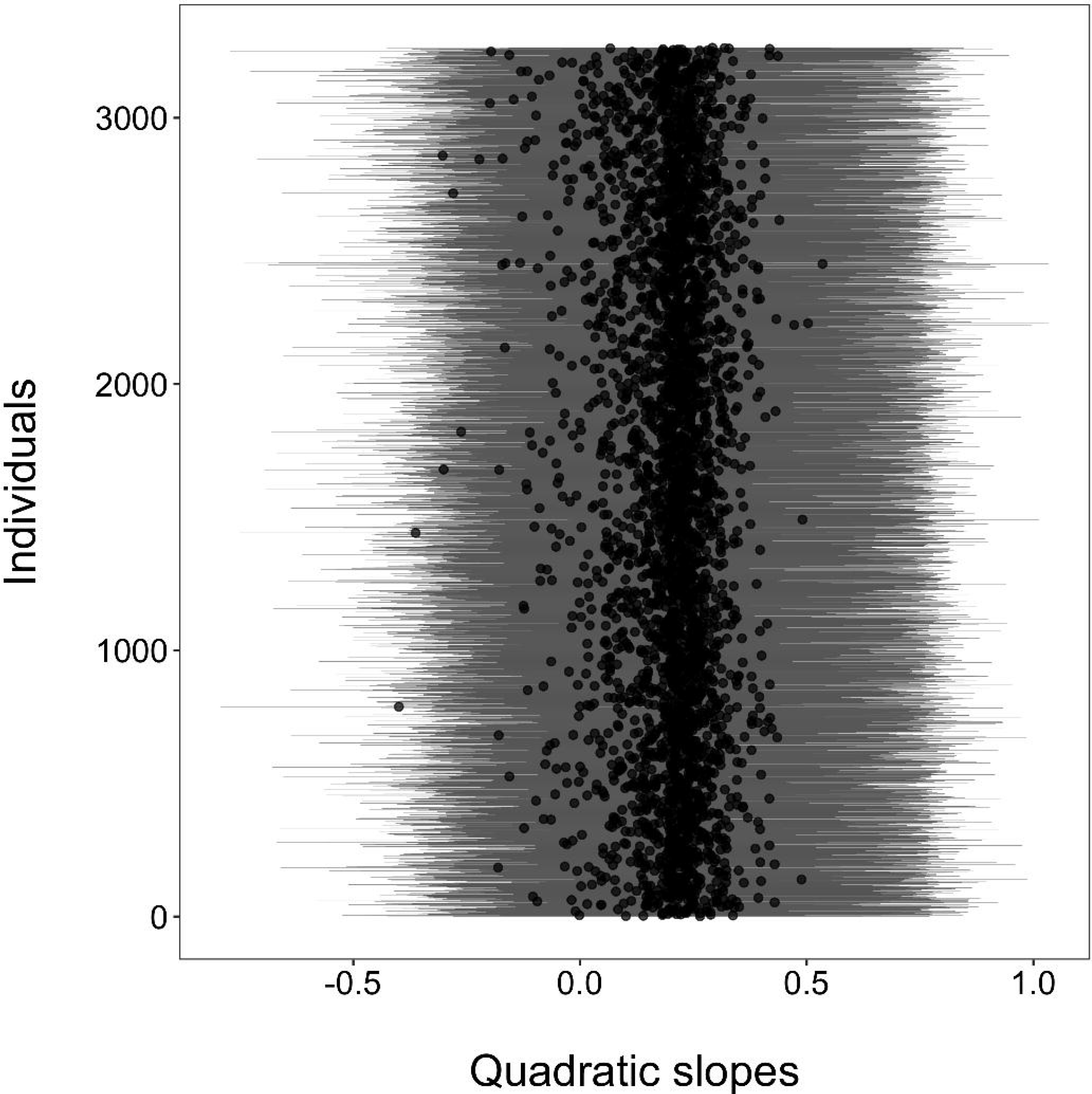
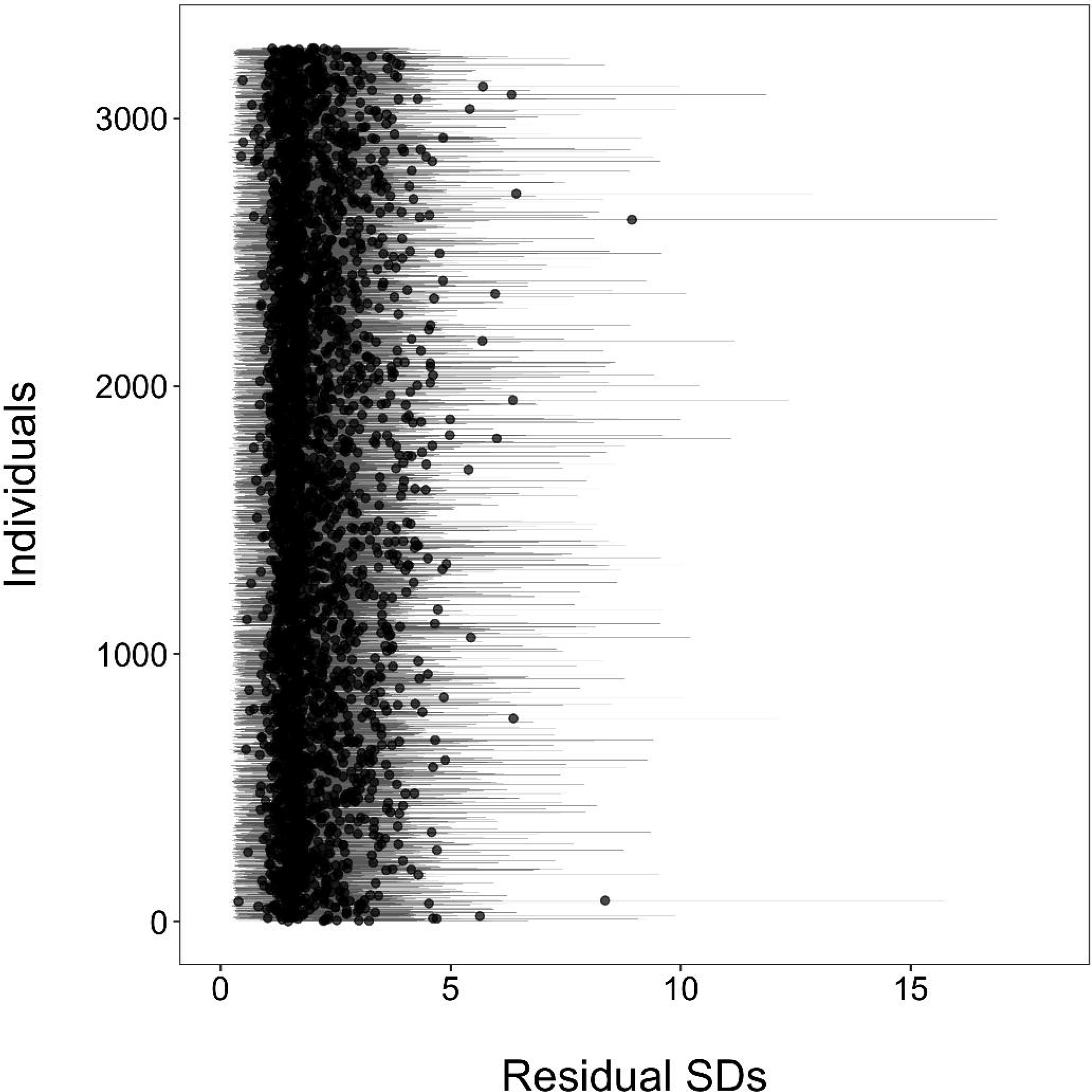
Posterior means (black dots) and 95% highest density intervals (grey vertical lines) for each dogs’ (a) intercept, (b) linear slope, (c) quadratic slope, and (d) residual SD parameter.

**Figure 4.**
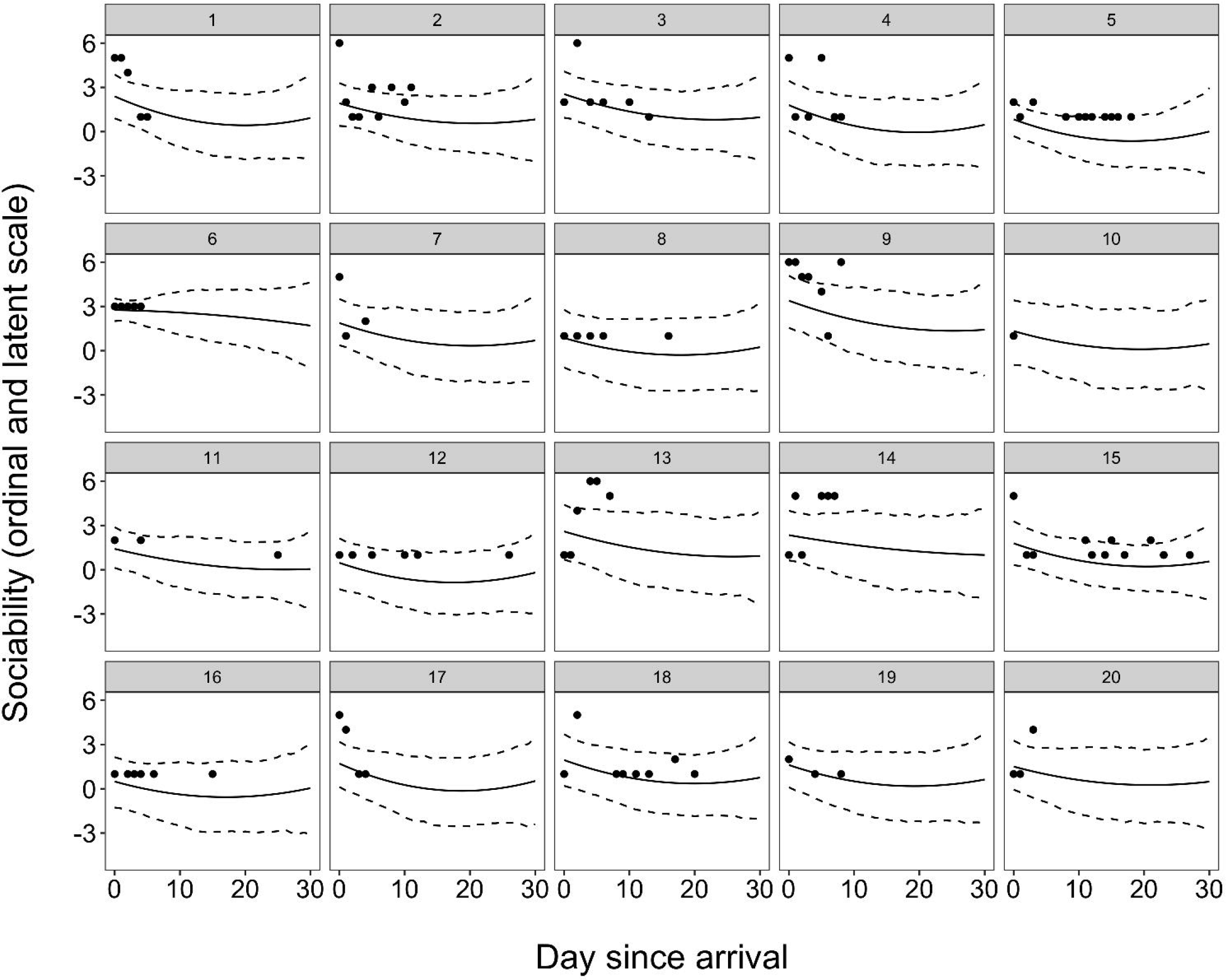
Predicted reaction norms (‘counterfactual’ plots) for twenty randomly-selected dogs. Black points show raw data on the ordinal scale (higher values indicate lower sociability), and solid and dashed lines illustrate posterior means and 95% highest density intervals. When data were sparse, there was increased uncertainty in model predictions. Due to hierarchical shrinkage, individual dogs' model predictions were pulled towards the group-level mean, particularly for those dogs showing higher behavioural codes (i.e. less sociable responses).

**Figure 5.**
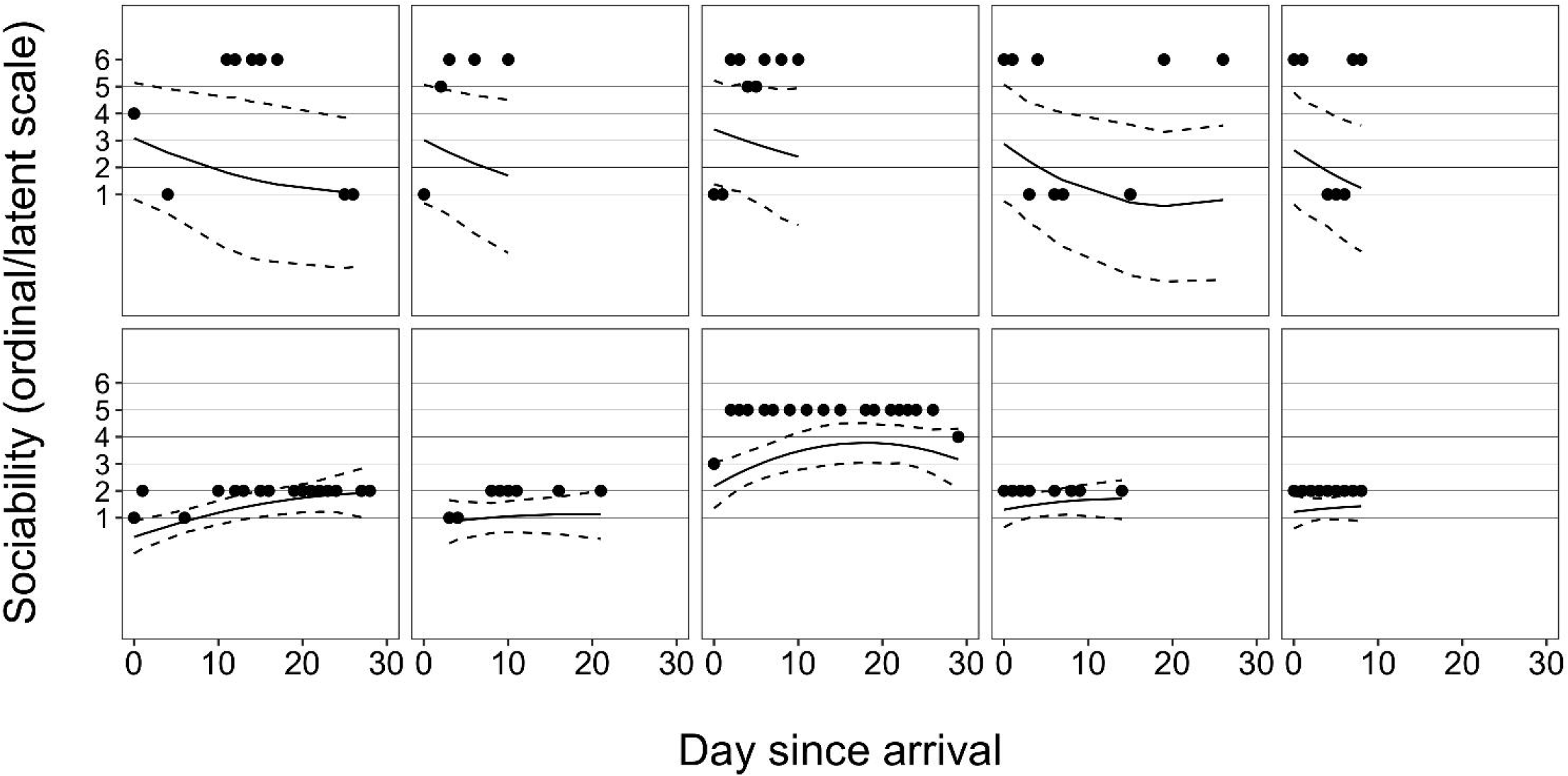
Reaction norms (posterior means = solid black lines; 95% highest density intervals = dashed black lines) for individuals with the five highest (top row) and five lowest (bottom row) residual SDs. Black points represent raw data on the ordinal scale (higher values indicating lower sociability).

Dog-varying intercepts positively correlated with linear slope parameters (*ρ* = 0.38, 95% HDI: 0.24, 0.50) and negatively correlated with quadratic slope parameters (*ρ* = −0.54, 95% HDI: −0.68, −0.39), and linear and quadratic slopes had a negative correlation (*ρ* = − 0.75, 95% HDI: −0.88, −0.59), indicating that less sociable individuals (with higher scores on the ordinal scale) had flatter reaction norms on average. Dog-varying residual SDs had a correlation with the intercept parameters of approximately zero (*ρ* = 0.00, 95% HDI: − 0.10, 0.10) but were negatively correlated with the linear slope parameters (*ρ* = −0.37, 95% HDI: −0.51, −0.22) and positively correlated with the quadratic slopes (*ρ* = 0.24, 95% HDI: 0.05, 0.42), indicating that dogs with greater residual SDs were predicted to change the most across days since arrival.

The ICC by day increased from arrival day (ICC = 0.22; 95% HDI: 0.16, 0.28) to day 8 (ICC = 0.33; 95% HDI: 0.28, 0.38) but changed little by day 15 (ICC = 0.32; 95% HDI: 0.27, 0.37). The cross-environmental correlation between days 0 and 8 was 0.79 (95% HDI: 0.70, 0.88), between days 0 and 15 was 0.51 (95% HDI: 0.35, 0.68), and between days 8 and 15 was 0.95 (95% HDI: 0.93, 0.97).

A one SD increase in the number of observations was associated with higher intercepts (*β* = 0.12; 95% HDI: 0.03, 0.21; see Supplementary Material Table S2) and higher residual SDs (*β* = 0.06, 95% HDI: 0.02, 0.10). Increasing age by one SD was associated with lower intercepts (*β* = −0.61, 95% HDI: −0.70, −0.51), steeper linear slopes (*β* = −0.20, 95% HDI: −0.27, −0.13), a stronger quadratic curve (*β* = 0.07, 95% HDI: 0.03, 0.12), and larger residual SDs (*β* = 0.05, 95% HDI: 0.01, 0.09). Increasing weight by one SD was associated with shallower quadratic curves (*β* = −0.05, 95% HDI: −0.09, −0.01). No credible effect of sex was observed on personality, plasticity or predictability. Gift dogs dogs (*β*_*diff*_= 0.28, 95% HDI: 0.04, 0.52) and stray dogs (*β*_*diff*_ = 0.33, 95% HDI: 0.15, 0.50), as well as steeper linear slopes (*β*_*diff*_ = −0.25, 95% HDI: −0.38, −0.13) and higher residual SDs than stray dogs (*β*_*diff*_ = 0.10, 95% HDI: 0.02, 0.18). Dogs at the large rehoming centre had steeper linear slopes (*β*_*diff*_ = −0.70, 95% HDI: −0.84, −0.56) and stronger quadratic curves (*β*_*diff*_ = 0.35, 95% HDI: 0.26, 0.45) than dogs at the medium rehoming centre, and lower intercept parameters (*β*_*diff*_ = − 0.30, 95% HDI: −0.50, −0.09) and steeper linear slopes (*β*_*diff*_ = −0.22, 95% HDI: −0.38, − 0.06) than dogs at the small rehoming centre. Compared to dogs at the small rehoming centre, dogs at the medium centre had lower intercepts (*β*_*diff*_ = −0.25, 95% HDI: −0.48, − 0.01), and shallower linear (*β*_*diff*_ = 0.48, 95% HDI: 0.30, 0.66) and quadratic slopes (*β*_*diff*_ = −0.34, 95% HDI: −0.46, −0.22). Dogs already neutered before arrival to the shelter had lower intercepts (*β*_*diff*_ = −0.54, 95% HDI: −1.07, −0.03) and lower residual SDs (*β*_*diff*_ = −0.53, 95% HDI: −0.85, −0.22) than dogs not neutered, but higher intercepts (*β*_*diff*_ = 0.20, 95% HDI: 0.03, 0.37) and higher residual SDs (*β*_*diff*_ = 0.10, 95% HDI: 0.02, 0.19) than those neutered whilst at the shelter. Unneutered dogs had higher intercepts (*β*_*diff*_ = 0.74, 95% HDI: 0.20, 1.26) and higher residual SDs (*β*_*diff*_ = 0.63, 95% HDI: 0.30, 0.92) than dogs neutered at the shelter.

## Discussion

This study applied the framework of behavioural reaction norms to quantify inter- and intra-individual differences in shelter dog behaviour during interactions with unfamiliar people. This is the first study to systematically analyse behavioural data from a longitudinal, observational assessment of shelter dogs. Dogs demonstrated substantial individual differences in personality, plasticity and predictability, which were not well described by simply investigating how dogs behaved on average. In particular, accounting for individual differences in predictability, or the short-term, day-to-day fluctuations in behaviour, resulted in significant improvement in model fit (Figure 1). Modelling dogs' longitudinal behaviour also demonstrated that behavioural repeatability increased with days since arrival (i.e. increasing proportion of variance explained by between-individual differences), particularly across the first week since arrival. Similarly, while individuals maintained rank-order differences in sociability across smaller periods (i.e. first 8 days), rank-order differences were only moderately maintained between arrival at the shelter and day 15. The results highlight the importance of adopting observational and longitudinal assessments of shelter dog behaviour, provide a method by which to analyse longitudinal data commensurate with other work in animal behaviour, and identify previously unconsidered behavioural measures that could be used to improve the predictive validity of behavioural assessments in dogs.

### Average behaviour

At the group-level, dogs' reactions to meeting unfamiliar people were predominantly coded as *Friendly* (Figure 2a), described as ‘Dog initiates interactions in an appropriate social manner’. Although this definition is broad, it represents a functional qualitative characterisation of behaviour suitable for the purposes of the shelter when coding behavioural interactions, and its generality may partly explain why it was the most prevalent category. The results are consistent with findings that behaviours indicative of poor welfare and/or difficulty of coping (e.g. aggression) are relatively infrequent even in the shelter environment [22, 26]. The change of behaviour across days since arrival was characterised by an increase in the *Friendly* code and a decrease in other behavioural codes (Figure 2a). Furthermore, the positive quadratic effect of day since arrival on sociability illustrates that the rate of behavioural change was not constant across days, being quickest earlier after arrival (Figure 2b). The range of behavioural change at the group-level was, nevertheless, still concentrated around the lowest behavioural codes, *Friendly* and *Excitable*.

Previous studies provide conflicting evidence regarding how shelter dogs adapt to the kennel environment over time, including behavioural and physiological profiles indicative of both positive and negative welfare [26]. Whereas some authors report decreases in the prevalence of some stress- and/or fear related behaviour with time [27, 49], others have reported either no change or an increase in behaviours indicative of poor welfare [17, 30]. Of relevance here, Kis *et al.* [17] found that aggression towards unknown people increased over the first two weeks of being at a shelter. In the current study, aggression was rare (Table 2), and the probability of ‘red codes’ (which included aggression) decreased with days at the shelter (Figure 3a). A salient difference is that Kis *et al.* [17] collected data using a standardised behavioural test consisting of a stranger engaging in a ‘threatening approach’ towards dogs. By contrast, we used a large data set of behavioural observations recorded after non-standardised, spontaneous interactions between dogs and unfamiliar people. In recording spontaneous interactions, the shelter aimed to elicit behaviour more representative of a dog’s typical behaviour outside of the shelter environment than would be seen in a standardised behavioural assessment. Previously, authors have noted that standardised behavioural assessments may induce stress and inflate the chances of dogs displaying aggression [29], emphasising the value of observational methods of assessment in shelters [24]. While such observational methods are less standardised, they may have greater ecological validity by giving results more representative of how dogs will behave outside of the shelter. Testing the predictive value of observational assessments on behaviour post-adoption is the focus of ongoing research.

### Individual-level variation

When behavioural data are aggregated across individuals, results may provide a poor representation of how individuals in a sample actually behaved. Here, we found heterogeneity in dog behaviour across days since arrival, even after taking into account a number of dog-level predictor variables that could explain inter-individual differences. Variation in individuals’ average behaviour across days (i.e. variation in dogs’ intercept estimates) illustrated that personality estimates spanned a range of behavioural codes, although model predictions mostly spanned the green codes (Figure 2b; Table 2). However, whilst there were many records to inform group-level estimates, there were considerably fewer records available for each individual, which resulted in large uncertainty of individual personality parameters (illustrated by wide 95% HDI bars in Figure 3a). Personality variation has been the primary focus of previous analyses of individual differences in dogs, often based on data collected at one time point and usually on a large number of behavioural variables consolidated into composite or latent variables (e.g. [50–52]). Our results highlight that ranking individuals on personality dimensions from few observations entails substantial uncertainty.

Certain studies on dog personality have explored how personality trait scores change across time periods, such as ontogeny (e.g. [53]) or time at a shelter (e.g. [17]). Such analyses assume, however, that individuals have similar degrees of change through time. If individuals differ in the magnitude or direction of change (i.e. degree of plasticity), group-level patterns of change may not capture important individual heterogeneity. In this study, most dogs were likely to show lower behavioural codes/more sociable responses across days since arrival, although the rate of linear and quadratic change differed among dogs. Indeed, some dogs showed a *decrease* in sociability through time (individuals with positive model estimates in Figure 3b), and while most dogs showed greater behavioural change early after arrival, others showed slower behavioural change early after arrival (individuals with negative model estimates in Figure 3c). As with estimates of personality, there was also large uncertainty of plasticity.

Part of the difficulty of estimating reaction norms for heterogeneous data is choosing a function that best describes behavioural change. We examined both linear and quadratic effects of day since arrival based on preliminary plots of the data, and their inclusion in the best fitting full model is supported by the lower WAIC value of alternative model 3, with both effects, compared to 4, with just the linear effect (Figure 1). Most studies are constrained to first-order polynomial reaction norms through time due to collecting data at only a few time points [6, 44]. However, the quadratic function was relatively easy to vary across individuals while maintaining interpretability of the results. More complex functions (e.g. regression splines) have the disadvantage of being less easily interpretable and higher-order polynomial functions may produce only crude representations of data-generating processes [33]. Nevertheless, by collecting data more intensely, the opportunities to model behavioural reaction norms beyond simple polynomial effects of time should improve. For instance, ecological momentary assessment studies in psychology point to possibilities for modelling behaviour as a dynamic system, such as with the use of vector-autoregressive models and dynamic network or factor models (e.g. [54, 55]). These models can also account for relationships between multiple dependent variables (e.g. multiple measures of sociability). Models of behavioural reaction norms, by contrast, have usually been applied to only one dependent variable operationally defined as reflecting the trait of interest, so methods to model multiple dependent variables through time concurrently will be an important advancement.

Personality and plasticity were correlated, with dogs with less sociable behaviour across days being less plastic. Previous studies have explored the relationship between how individuals behave on average and their degree of behavioural change. David *et al.* [56] found that male golden hamsters (*Mesocricetus auratus*) showing high levels of aggression in a social intruder paradigm were slower in adapting to a delayed-reward paradigm. In practice, the relationship between personality and plasticity is probably context dependent. Betini and Norris [57] found, for instance, that more aggressive male tree swallows (*Tachycineta bicolor*) during nest defence were more plastic in response to variation in temperature, but that plasticity was only advantageous for nonaggressive males and no relationship was present between personality and plasticity in females. The correlation between personality and plasticity indicates a ‘fanning out’ shape of the reaction norms through time (Figure 2b). Consequently, behavioural repeatability or the amount of variance explained by between-individual differences increased as a function of day, but only after the first week after arrival. The ‘cross-environmental’ correlation, moreover, indicated that the most sociable dogs on arrival day were not necessarily the most sociable on later days at the shelter. In particular, the correlation between sociability scores on arrival day and day 15 was only moderate, supporting Brommer [44] that the rank-ordering of trait scores is not always reliable. By contrast, the cross-environmental correlations between days 0 and 8, and between days 8 and 15, were much stronger. These results suggest that shelters using standardised behavioural assessments would benefit from administering such tests as late as possible after dogs arrive.

Of particular interest was predictability or the variation in dogs’ residual SDs. Studies of dog personality generally treat behaviour as probabilistic, implying recognition that residual intra-individual behaviour is not completely stable, and authors have posited that dogs may vary in their behavioural consistency (e.g. [13]). Yet, this is the first study to quantify individual differences in predictability in dogs. Modelling residual SDs for each dog resulted in a model with markedly better out-of-sample predictive accuracy (Figure 1). The coefficient of variation for predictability was 0.64 (95% HDI: 0.58, 0.70), which is high compared to other studies in animal behaviour. For instance, Mitchell *et al.* [6] reported a value of 0.43 (95% HDI: 0.36, 0.53) in spontaneous activity measurements of male guppies (*Poecilia reticulata*). Variation in predictability also supports the hypothesis that dogs have varying levels of behavioural consistency. It is important to note, however, that interactions with unfamiliar people at the shelter were likely more heterogeneous than behavioural measures from standardised tests or laboratory environments, which may contribute to greater individual variation in predictability. Moreover, the behavioural data analysed here may have contained more measurement error than data from more standardised environments.

Although shelter employees demonstrated significant inter-rater reliability in video coding sessions, the average proportion of shelter employees who selected the correct behavioural code to describe behaviours seen in videos was modest (66%), while 78% chose a video in the correct colour category (green, amber or red). Indeed, only 55% of employees identified the *Reacts to people aggressive* behaviour as a red code, with the remaining employees identifying it as the amber category code *Reacts to people non-aggressive*. As discussed by Goold and Newberry [35], employees were likely to mistake examples of aggression for non-aggression, but not the other way around. In the current study, this would have increased the percentage of lower category codes (describing greater sociability). Due to lower standardisation of the observational contexts at the shelter than in formal behavioural testing, it was important to evaluate the reliability and validity of the behavioural records. Defining acceptable standards of reliability and validity is, however, non-trivial and we could not find measures of reliability or validity in any previous studies investigating predictability in animals for comparison.

Dogs with higher residual SDs demonstrated steeper linear slopes and greater quadratic curves, indicating that greater plasticity was associated with lower predictability. The costs of plasticity are believed to include greater phenotypic instability, in particular developmental instability [11, 58]. Since more plastic individuals are more responsive to environmental perturbation, a limitation of plasticity may be greater phenotypic fluctuation on finer time scales. However, lower predictability may also confer a benefit to individuals precisely because they are less predictable to con- and hetero-specifics. For instance, Highcock and Carter [59] reported that predictability in behaviour decreases under predation risk in Namibian rock agamas (*Agama planiceps*). No correlation was found here between personality and predictability, similar to findings of Biro and Adriaenssens [2] in mosquitofish (*Gambusia holbrooki*), although correlations were found in agamas [59] and guppies [6]. It is possible that correlations between personality and predictability depend upon the specific aspects of personality under investigation.

### Predictors of individual variation

Finally, we found associations between certain predictor variables and personality, plasticity and predictability (Supplementary Material Table S2). Our primary reason for including these predictor variables was to obtain more accurate estimates of personality, plasticity and predictability, and we remain cautious about *a posteriori* interpretations of their effects, especially since the theory underlying why individuals may, for example, demonstrate differences in predictability is in its infancy [8]. The reproducibility of a number of the results would, nevertheless, be interesting to confirm in future research. In particular, understanding factors affecting intra-individual change is important given that many personality assessments are used to predict an individual’s future behaviour, rather than understand inter-individual differences. Here, increasing age was associated with greater plasticity (linear and quadratic change) and lower predictability, although some of the parameters’ 95% HDIs were close to zero, indicative of small effects. In great tits (*Parus major*) conversely, plasticity decreased with age [60], whilst in humans, intra-individual variability in reaction times increased with age [61]. Moreover, non-neutered dogs showed lower predictability than neutered dogs, and dogs entering the shelter as gifts (relinquished by their owners) had lower predictability estimates than stray dogs (dogs brought in by local authorities or members of the public after being found without their owners). These results can be used to formulate specific hypotheses about behavioural variation.

## Conclusion

We applied the framework of behavioural reactions norms to data from a longitudinal and observational shelter dog behavioural assessment, quantifying inter- and intra-individual behavioural variation in dogs’ interactions with unfamiliar people. Overall, shelter dogs were sociable with unfamiliar people and sociability continued to increase with days since arrival to the shelter. At the same time, dogs showed individual differences in personality, plasticity and predictability. Accounting for all of these components substantially improved model fit, particularly the inclusion of predictability, which suggests that individual differences in day-to-day behavioural variation represent an important, yet largely unstudied, component of dog behaviour. Our results also highlight the uncertainty of making predictions about shelter dog behaviour, particularly when the number of behavioural observations is low. For shelters conducting standardised behavioural assessments, assessments are likely best carried out as late as possible, given that rank-order differences between individuals on arrival and at day 15 were only moderately related. In conclusion, this study supports moving towards observational and longitudinal assessments of shelter dog behaviour, has demonstrated a Bayesian method by which to analyse longitudinal data on dog behaviour, and suggests that the predictive validity of behavioural assessments in dogs could be improved by systematically accounting for both inter- and intra-individual variation.

## Ethics statement

Full permission to use the data in this article was provided by Battersea Dogs and Cats Home.

## Data accessibility

The data, R code and Stan model code to run the analyses and produce the results and figures in this article are available on Github: https://github.com/ConorGoold/GooldNewberry_modelling_shelter_dog_behaviour

## Competing interests

We declare no competing interests.

## Author contributions

CG and RCN conceptualised the study. CG obtained the data, conducted the statistical analyses and drafted the initial manuscript. CG and RCN revised the manuscript and wrote the final version.

## Acknowledgements

The authors are especially grateful to Battersea Dogs and Cats Home for providing the data on their behavioural assessment.

## Funding statement

CG and RCN are employed by the Norwegian University of Life Sciences. No additional funding was required for this study.

